# Extended lifespan in female *Drosophila melanogaster* through late-life caloric restriction

**DOI:** 10.1101/2023.07.13.548896

**Authors:** Michael Li, Jacob Macro, Billy J. Huggins, Kali Meadows, Dushyant Mishra, Dominique Martin, Blanka Rogina

## Abstract

Calorie restriction has many beneficial effects on healthspan and lifespan in a variety of species. However, how late in life application of caloric restriction can extend fly life is not clear. Here we show that late-life calorie restriction increases lifespan in female *Drosophila melanogaster* aged on a high calorie diet. This shift results in rapid decrease in mortality rate and extends fly lifespan. In contrast, shifting female flies from a low to a high calorie diet leads to a rapid increase in mortality and shorter lifespan. These changes are mediated by immediate metabolic and physiological adaptations. One of such adaptation is rapid adjustment in egg production, with flies directing excess energy towards egg production when shifted to a high diet, or away from reproduction in females shifted to low caloric diet. However, lifelong female fecundity reveals no associated fitness cost due to CR when flies are shifted to a high calorie diet. In view of high conservation of the beneficial effects of CR on physiology and lifespan in a wide variety of organisms, including humans, our findings could provide valuable insight into CR applications that could provide health benefits later in life.

## Introduction

Calorie restriction (CR) is a robust intervention known to extend lifespan across many model organisms (1–3). This nutritional intervention has been well studied in multiple contexts, identifying countless factors that can influence an organism’s response to CR, including, diet composition, caloric content, time of application, genetic background, sex, among others (1,2,4–7). Despite this, much remains to be elucidated regarding underlying mechanisms and its potential utility in extending human health. One key question that is essential to address in this regard is whether organisms can still benefit from CR when applied late in life.

One model system that has been extensively utilized in order to investigate CR is the fruit fly, *Drosophila melanogaster* (8). Using this system, many studies have found that CR is beneficial in extending fly lifespan through various mechanisms (2,7,8). Interestingly, these mechanisms are not unique to flies, as CR exerts similar effects across several model systems, including, mice, worms and nonhuman primates (2,9,10). Thus, information learned from flies may be translatable to humans, as the mechanisms driving these responses seem to be highly conserved (11,12).

Previous work has demonstrated that flies can benefit from switching from a high to a low calorie diet, resulting in lifespan extension despite a history on a rich diet (13–15). However, the opposite was also true, with flies switched from a low to a high calorie diet demonstrating an immediate increase in risk of death. In some studies, this risk was even higher than that of flies on a lifelong, high calorie diet(14). In this particular study, the authors also reported restricted flies to be less fecund when returned to normal conditions(14). This finding suggested there may be a hidden cost associated with CR, challenging the view that restriction is a stimulus that cues an organism to invest in somatic maintenance. However, the hypothesis that CR imposes a hidden cost was later contested by data in an outbred population of female flies, showing CR did not result in a fitness cost that hindered fecundity upon refeeding(15). Such differences in experimental results could be due to various reasons, including, differing experimental designs and/or genetic background, as both can influence response to CR (16).

Here, we examined if wild type, female *Canton-S* (*CS*) flies could benefit from application of calorie restriction late in life. We also examined the effects of shifting diets with different caloric content on fly metabolic and physiological adaptation. Rather than shifting flies from standard to restricted diets, as has been done previously, we shifted flies from a high to a low calorie diet (and vice versa) in order to increase clinical relevance. We reasoned that since most humans with metabolic disorders eat above the daily recommended allowance, shifting flies to and from a high calorie diet would better recapitulate human response to CR (17). Due to the increase in age-associated incidence of metabolic syndrome, maintaining a better understanding of age-dependent application of CR is vital.

## Methods

### Fly strains, maintenance and diet

We used the wild-type *Canton-S (CS)* line obtained from the Bloomington Stock Center (Stock number 1). *CS* flies were reared on food containing 25 mg/mL tetracycline for 3 generations to eliminate *Wolbachia*. This treatment was followed by growing flies for generations in tetracycline-free food. Standard laboratory corn media was used to grow *CS* fly grandparents and parents from which cross was set to for F1 fly collections(18). The time each group of flies stay in one vial starting with grandparents was the same to avoid any effects of larval density on longevity. Each cross for grandparents and parents was done by crossing 10 virgin female and 9 male flies who were 4–10 days old. The flies were passed to a new vial with corn media every 2 days. F1 progeny was collected every 24 hours. 25 male and 25 female flies were kept on a low (L) or high (H) food per experimental design. Flies were maintained in a humidified temperature-controlled environmental chamber at 25°C (Percival Scientific) on a 12-hour light:dark cycle with light on at 6 AM. Standard laboratory corn media was used for setting the crosses, while experimental diet is marked as low, L = 0.5N and high H = 3.0N (18). The two food caloric levels are standardized as 1.0N being the food that has 100 g/L of sucrose (MP Biomedicals, Inc), 100 g/L of brewer’s yeast (MP Biomedicals, Inc) and 20 g/L of agar (8,18,19). Food preparation was described previously (18).

### Lifespan Studies

Lifespan studies were performed using 10 to 12 vials of 25 male and 25 female flies placed in each vial. Flies were collected within 24 hours following eclosion and maintained in plastic vials containing high or low-calorie diet, and maintained as described above. While males were aged together with female flies, here we present data for only female flies, while male findings are presented elsewhere (20). The number of flies in each survivorship study is listed in Tables 1, 2 and Supplemental Tables 1,2. Two different shifting experiments were performed. In experiment 1: Two groups of *CS* flies were aged on a high-calorie (H) or low-diet (L) for their whole lifespan. Additional three groups of flies aged on L, and three groups aged on H diet, were shifted to opposite food at ages 20, 50, or 60 days. Flies were passed every day from day 1 and the number of dead flies were counted. Experiment 2: Two groups of *CS* flies were aged their whole lifespan on a H or L diets, and five additional groups that began on H from birth and were moved to L at either 10, 20, 30, 40 or 50 days. Five additional groups of flies were kept on L diet from birth and shifted from L to H at either 10, 20, 30, 40 or 50 days. They were passed every 2 days up to age 10 days and every day after that and the number of dead flies were counted.

**Table 1:**
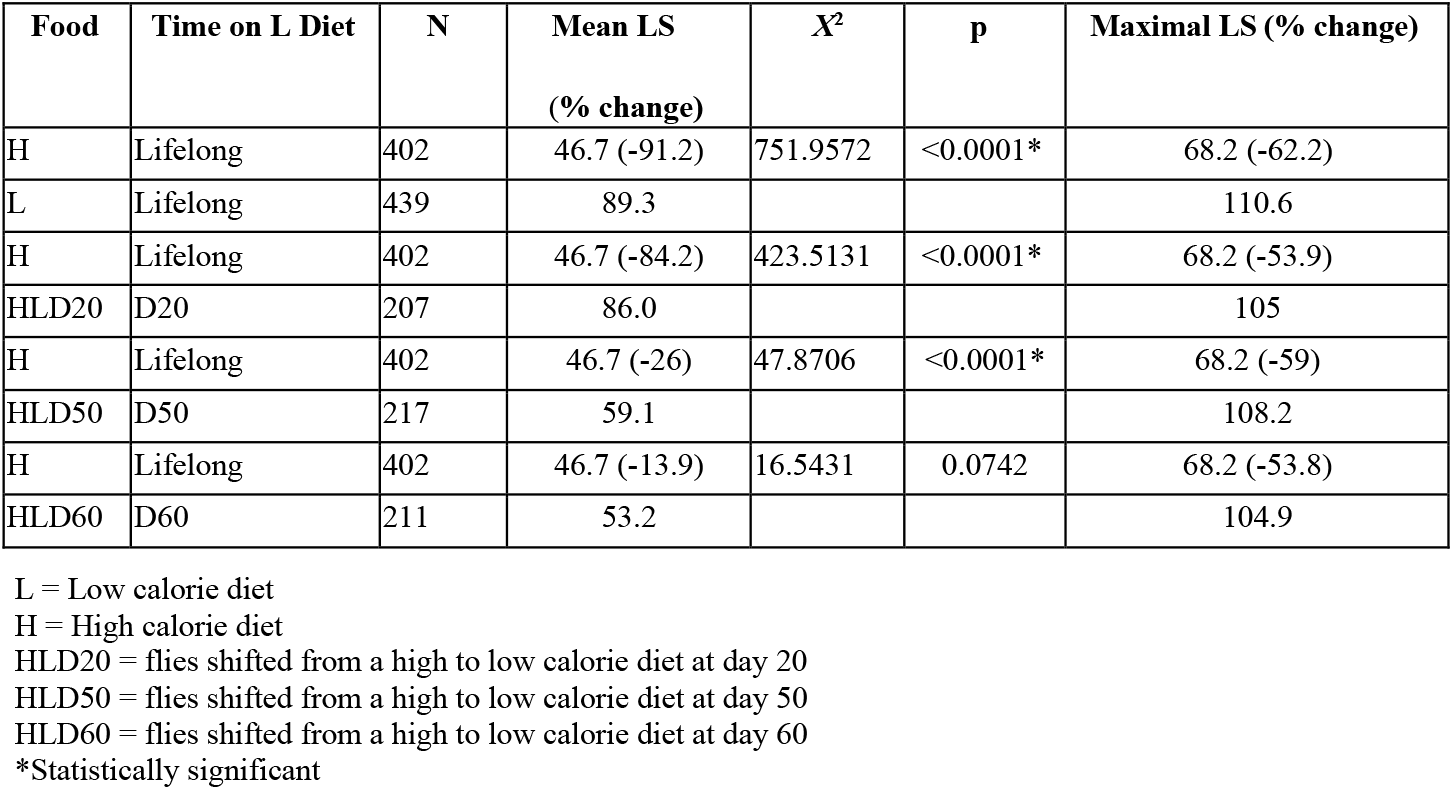
Effects of shifting *Canton-S* female flies from a high (H) to a low (L) calorie diet on longevity compared to longevity of flies on lifelong high calorie diet.

| Food | Time on L Diet | N | Mean LS<br>(% change) | $X^2$ | p | Maximal LS (% change) |
| --- | --- | --- | --- | --- | --- | --- |
| H | Lifelong | 402 | 46.7 (-91.2) | 751.9572 | <0.0001* | 68.2 (-62.2) |
| L | Lifelong | 439 | 89.3 |  |  | 110.6 |
| H | Lifelong | 402 | 46.7 (-84.2) | 423.5131 | <0.0001* | 68.2 (-53.9) |
| HLD20 | D20 | 207 | 86.0 |  |  | 105 |
| H | Lifelong | 402 | 46.7 (-26) | 47.8706 | <0.0001* | 68.2 (-59) |
| HLD50 | D50 | 217 | 59.1 |  |  | 108.2 |
| H | Lifelong | 402 | 46.7 (-13.9) | 16.5431 | 0.0742 | 68.2 (-53.8) |
| HLD60 | D60 | 211 | 53.2 |  |  | 104.9 |

L = Low calorie diet
H = High calorie diet
HLD20 = flies shifted from a high to low calorie diet at day 20
HLD50 = flies shifted from a high to low calorie diet at day 50
HLD60 = flies shifted from a high to low calorie diet at day 60
\*Statistically significant

**Table 2:** Effects of shifting *Canton-S* female flies from a low (L) to a high (H) calorie diet on longevity compared to longevity of flies on lifelong low calorie diet.

| Food | Time on L Diet | N | Mean LS<br>(% change) | $\chi^2$ | p | Maximal LS<br>(% change) |
| --- | --- | --- | --- | --- | --- | --- |
| L | Lifelong | 439 | 89.3 (-48) | 751.9572 | <0.0001* | 110.6 (-38) |
| H | Lifelong | 402 | 46.7 |  |  | 68.2 |
| L | Lifelong | 439 | 89.3 (-44) | 565.7714 | <0.0001* | 110.6 (-31) |
| LHD20 | D20 | 207 | 49.8 |  |  | 76.3 |
| L | Lifelong | 439 | 89.3 (-35) | 526.5062 | <0.0001* | 110.6 (-30) |
| LHD50 | D50 | 226 | 58.0 |  |  | 77.9 |
| L | Lifelong | 439 | 89.3 (-31) | 454.1635 | <0.0001* | 110.6 (-26) |
| LHD60 | D60 | 218 | 62.0 |  |  | 82.2 |

L=Low calorie diet
H=High calorie diet
LHD20= flies shifted from a low to high calorie diet at day 20
LHD50= flies shifted from a low to high calorie diet at day 50
LHD60= flies shifted from a low to high calorie diet at day 60
\*Statistically significant

### Fecundity

Life-long fecundity was determined from daily counts of eggs laid by individual females from 20 replicate vials of the *CS* flies in each experiment: two groups of flies kept on L and H calorie diet, and another two groups of flies were shifted from L to H or H to L at 10 or 50 days. Each *CS* females was placed along with one male flies per vial. Each day, they were passed into new vials, and the number of eggs present was recorded (21). Number of eggs is listed in Table 3.

**Table 3:** Effects of shifting *Canton-S* female flies from a high to a low or from a low to a high calorie diet on fecundity.

| **Diet** | **Time on Diet** | **Lifelong**<br>**N eggs** | **SEM** | **6-10D** | **SEM** | **11-15D** | **SEM** |
| --- | --- | --- | --- | --- | --- | --- | --- |
| H | Lifelong | 906.5 | 79.0 | 221 | 19.2 | 206 | 19.8 |
| HLD10 | 0-10 H, 10-rest L | 884.80 | 68.9 | 243 | 23.2 | 114 | 9.0 |
| LHD10 | 0-10 L, 10-rest H | 668.65 | 86.2 | 70 | 4.8 | 172 | 21.1 |
| L | Lifelong | 593.4 | 31.6 | 58 | 3.2 | 77 | 2.0 |

Average number of egg/fly for *Canton-S* female flies kept on a high (H) or a low (L) calorie diet lifelong, or shifted from a H to a L, or L to H at day 10. The number of average eggs/fly listed are laid during female lifetime, during 6-10, or 11-15 day periods. Each *CS* females was placed along with one male flies per vial on day 0. SEM= of the average eggs laid during listed periods. There were 20 females in each experiment. N=number of eggs.
L=Low calorie diet
H=High calorie diet
HLD10= flies shifted from a high to low calorie diet at day 10
LHD10= flies shifted from a low to high calorie diet at day 10

### Biochemistry

*Canton-S (CS)* flies were collected and aged as described above on L or H diets. At ages 20 and 50, subgroups of flies were transferred to the opposite diet. Analysis was done on flies aged their whole life on L or H at age 20 or 50 days. Groups of flies transferred to opposite diet at 20 days were used for analysis at ages 21, 22, and 25 days. Similarly, flies that were transferred at 50 days to the opposite diet were used for analysis at ages 51, 52, and 55 days. Flies were sorted on CO_2_ by sex. 3 biological replicates of 10 female flies, were anesthetized on CO_2_, weighed, homogenized in homogenization buffer and tube was spun down at 2000 rpm for 2 minutes at 4°C. 25 μl of each homogenate was aliquoted into three wells of each of six 96-well plates. The plates were kept on dry ice allowing homogenates to be frozen immediately and kept at -80°C until quantification. Before quantification, the plates were left to warm up to room temperature for 15 minutes. For glucose, Peroxidase-Glucose oxidase (PGO) enzyme plus color reagent was added to each well (Glucose Assay Kit: Sigma GAG020, PGO Sigma P7119), the plate was incubated at 37°C and the optic density was read at 450 nm using a Tecan Spark Microplate Reader. For glycogen, the procedure was the same as glucose except amyloglucosidase (Amyloglucosidase from *Aspergillus niger*, Sigma 10115) was added to each well in addition to the other enzyme. For trehalose, the procedure was the same as glucose except the samples were incubated with trehalase at 37°C before adding PGO (Trehalase from porcine kidney, Sigma T8778). Protein was determined using Total Protein Kit, Micro Lowry, Peterson’s Modification (Sigma TP0300). The plate was incubated at room temperature and was read at 750 nm. The triglycerides were determined enzymatically using Serum Triglyceride Determination Kit (Sigma TR0100; Glycerol Standard Solution Sigma G7793)(22).

### Statistical analysis

Egg production data from period between 6-10 and 11-15 days were analyzed separately using the Kruskal-Wallis test. Post-hoc analysis was conducted using Dunn’s test, correcting for multiple comparisons using GraphPad Prism 9.4.1, Results represent SEM. There were 20 female flies in each experiment. p: 0.033 (*), 0.002 (**), 0.0002 (***), <0.0001 (****).

Biochemistry data from 20- and 50-day-old flies were analyzed separately using two-way ANOVA. Post-hoc analysis was conducted using Tukey’s test, correcting for multiple comparisons using GraphPad Prism 9.4.1. Results represent means ± SE of 3 biological replicates containing 30 flies per replicate. p: 0.033 (*), 0.002 (**), 0.0002 (***), <0.0001 (****). Longevity data were censored for early mortality (1 - 10 Days) and analyzed by log-rank tests using the JMP16 program.

## Results

### Calorie restriction increases lifespan of female flies when applied late in life

In order to probe how late in life *CS* female flies aged on a high calorie diet can benefit from CR, we performed a series of survivorship studies in which flies were shifted from a high to a low calorie diet at various time points. In our initial survivorship studies, flies were shifted at days 10, 50, and 60, as indicated in Figure 1A. As expected, flies raised on a low calorie diet lived significantly longer than those raised on a high calorie diet (Figure 1B, Table 1, p<0.0001). Strikingly, flies shifted from a high to a low calorie diet on days 20 and 50 displayed significant lifespan extension (Figure 1B, Table 1, p<0.0001). The survivorship curve of flies shifted on day 20 almost overlaps with survivorship of flies on low calorie diet. When flies were shifted to a low calorie diet at the age of 50 days, the response was dramatic, with a 59% increase in maximal lifespan compared to flies on a lifelong, high calorie diet (Table 1). When shifted at day 60, lifespan was not significantly different from flies on a lifelong, high calorie diet (Figure 1B, Table 1, p=0.07), but the remaining, old flies still responded to a low calorie diet with a 53.8% increase in maximal lifespan. Both day 50 and day 60 shifts lead to dramatic changes in survivorship, which becomes almost flat after the shift and then follows a similar trajectory of survivorship of flies on a low calorie diet at about 100 days of age. In a second set of experiments performed, an immediate and profound increase in survivorship was also observed when flies were shifted from a high to a low diet starting at 10 days of age all the way to 50 days of age (Figure 2A). Again, flies shifted to a low calorie diet gained a significant increase in survivorship compared to flies on high calorie diet (Figure 2A, Supplemental Tables 1,2). *Together, these data suggest that female flies aged on a high calorie diet can benefit from CR, even when implemented at an old age*.

**Fig. 1.**
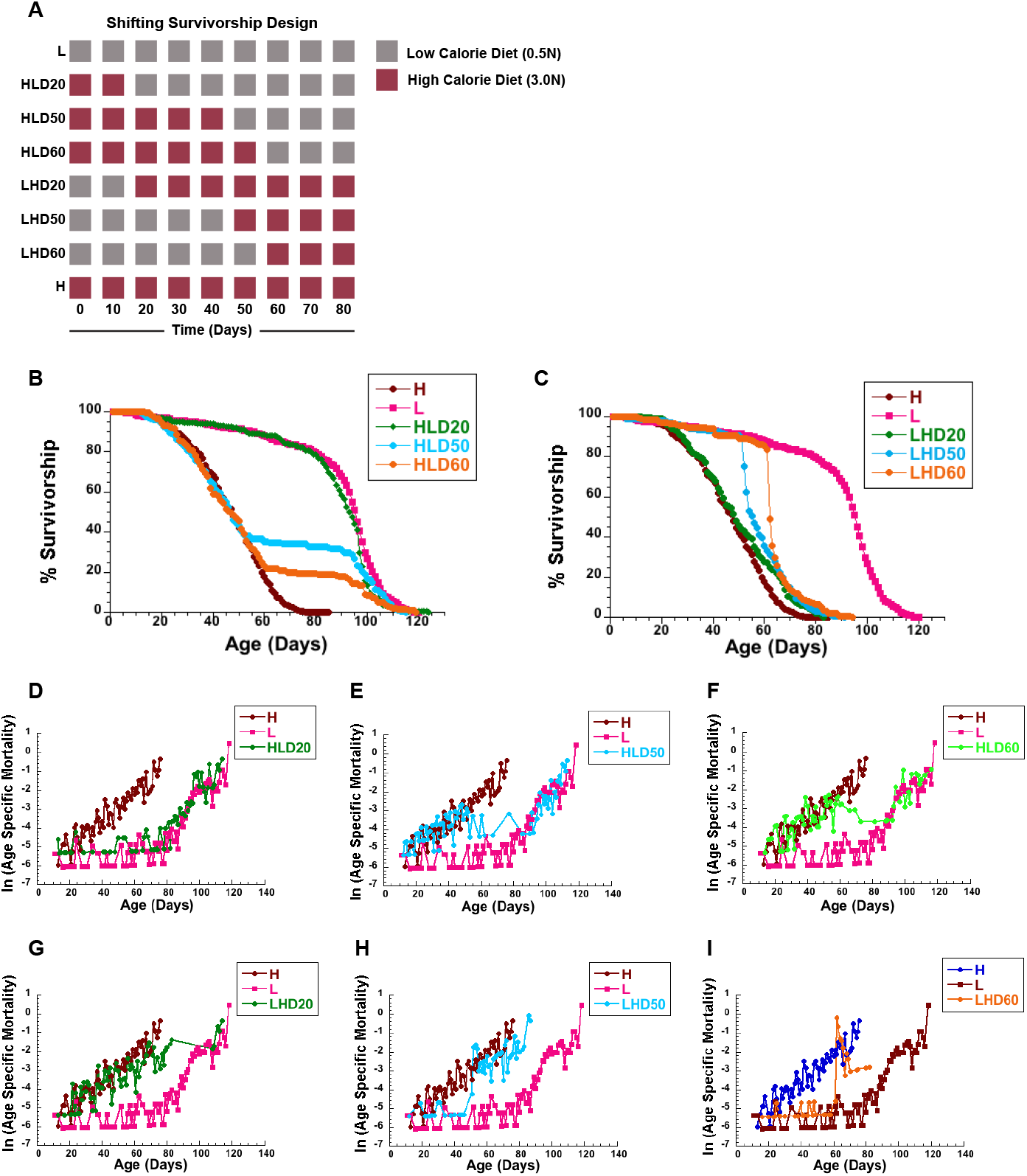
Shifting female flies to diet with different caloric content effects lifespan and mortality: **A** Schematic diagram of the experimental design: a group of female flies were aged on a low calorie (L), or a high calorie diet (H). Additional three groups of flies were shifted from H to L diet at day 20 (HLD20), 50 (HLD50), or 60 (HLD60), and another three from L to H diet on day 20 (LHD20), 50 (LHD50), or 60 (LHD60). **B-I** Survivorships (**B**,**C**) and mortality rates (**D-I**) of female flies shifted from H to L at day 20 (HLD20), day 50 (HDL50), or day 60 (HLD60) (**B**,**D-F**) or from L to H diet on day 20 (LHD20), day 50 (LHD50), or day 60 (LHD60) (**C**,**G-I**). Number of flies: L=439, H=402, HLD20=207, HLD50=217, HLD60=211, LHD20=207, LHD50=226, LHD60=218. Survivorship curves and mortality rate were analyzed by long-rank test JMP16 program.

**Fig. 2.**
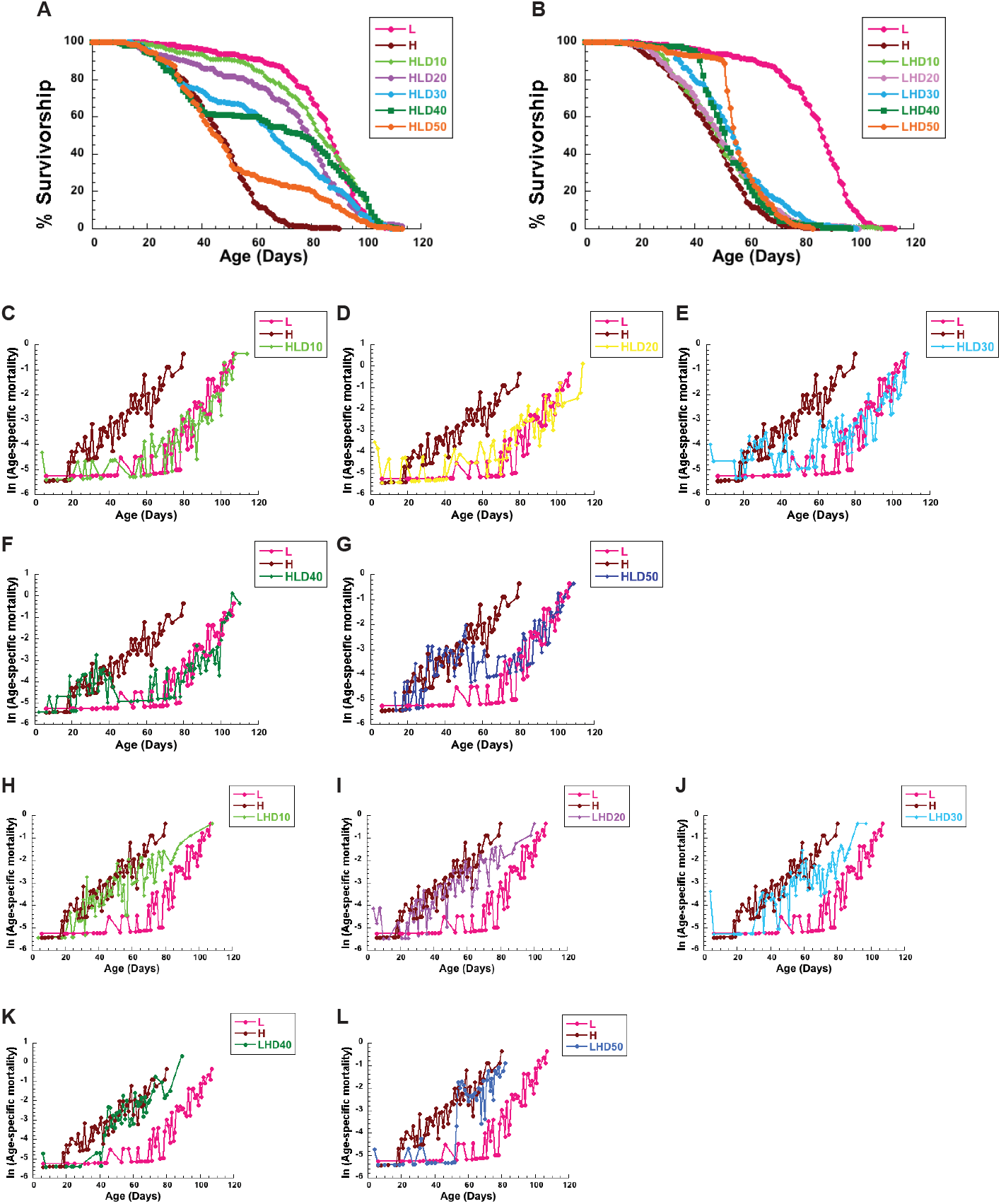
Shifting diets at different time during aging has immediate effects on female lifespan and mortality: Survivorships (**A**,**B**) and mortality rates (**C-L**) of female flies shifted from a high (H) to a low (L) calorie diet at day 10 (HLD10), day 20 (HLD20), day 30 (HLD30), day 40 (HLD40) or day 50 (HDL50) (**C-G**) or from L to H diet on day 10 (LHD10), day 20 (LHD20), day 30 (LHD30), day 40 (LHD40) or day 50 (LHD50) (H-L). Number of flies: L=194, H=227, HLD10=219, HLD20=225, HLD30=211, HLD40=222, HLD50=229, LHD10=231, LHD20=242, LHD30=199, LHD40=228, LHD50=224. Survivorships curves and mortality rate were analyzed by long-rank test JMP16 program.

To further examine the effects of late-life caloric shifting on female flies, we also performed opposite experiments in which flies were started on a low calorie diet, and then shifted to a high calorie diet at days 10, 50, and 60, as indicated in Figure 1A. In contrast, shifting from a low to a high calorie diet dramatically shortened female lifespan (Figure 1C, Table 2, p<0.0001). Interestingly, the shortening in lifespan was proportional to the age of the flies when shifted, with flies shifted at 60 days exhibiting a smallest, negative effects on lifespan. For example, when shifted to a high calorie diet at day 20, there was a 44% decrease in median lifespan compared to flies on a lifelong, low calorie diet (Table 2). However, when shifted at day 60, there was a 31% decrease in median lifespan between shifted flies and those on a lifelong, high calorie diet (Table 2). *Together, our data suggests that high calorie diet is detrimental to fly lifespan at all ages, despite previous exposure to a low calorie diet, but that lifespan is immediately responsive to caloric shifts*.

### Calorie restriction alters age-specific mortality

One advantage of performing survivorship experiments with a high sample size is the ability to calculate age-specific mortality rates. Such calculations can allow for comparisons of death vulnerability at different ages and can provide insight into the reasons for a given intervention’s effect (13). In this case, we were interested in the effects of age and caloric content on mortality risk in female flies. We first calculated age-specific mortality rates for flies that underwent shifts from a high to a low calorie diet. Under these conditions, flies responded with an immediate decrease in risk of death upon the switch, with rates quickly mirroring flies raised on a lifelong, low calorie diet (Figure 1D-F). This was true at each age tested, including 60 days, suggesting that low calorie diets decrease risk of death even after a history of high caloric diets. Importantly, the slope of the mortality trajectory was similar between switched flies and those on a lifelong, low calorie diet, suggesting a decrease in short term risk of death rather than a decrease in the accumulation of age-related damage when switched (13).

Considering previous evidence that flies on a restricted diet respond negatively when switched to fully fed conditions (14,15) we wanted to examine whether low calorie diets at the start of life would be protective or detrimental to flies when switched to a high calorie diet later in life. To do this, we also calculated age-specific mortality rates of flies that were shifted from a low to a high calorie diet. Our previous study in males indicated that flies gain an instantaneous increase in mortality risk when switched to a high calorie diet, one that is higher than that of flies raised on a constant, high calorie diet (20). This was not necessarily true of females. While there was an increase in risk at all ages tested when switched, younger flies (Day 20) seemed to be more resistant to the harmful effects of a high calorie diet than older flies (Day 50 and 60). At day 20, flies showed a mortality risk that was no higher than those on a lifelong, high calorie diet. As these flies aged, their risk of dying was lower than flies on a high calorie diet, resulting in a longer lifespan than flies aged on a lifelong, high calorie diet (Figure 1G). At days 50 and 60, there was a transient increase in risk that was higher than flies raised on a high calorie diet, but as flies in this group aged, they showed a lower risk of mortality than those on a high calorie diet, again, resulting in a longer lifespan than flies on constant, high calorie diet (Figure 1H and I, Supplemental Table 3). To further appreciate the increase in acute risk of death that occurs after switching from a low to high calorie diet at an old age, we evaluated survivorship immediately after the shift occurred (Supplemental Figure 1A). When evaluating survivorship in this way, it became immediately obvious that survivorship decreased rapidly immediately following the shift, at a rate that is faster than flies on a lifelong, high calorie diet, which corresponded with the increase in mortality around day 50 (Supplemental Figure 1A and B). However, by day 55, survivorship then proceeded at the same rate in flies switched from low to high as those on a high calorie diet (Supplemental Figure 1A). This effect was further exacerbated when the flies were switched at day 60 (Supplemental Figure 1C and D). This result was even more obvious in our second survivorship study, which showed low to high shifted flies have an intermediate increase in mortality rate that is higher than flies on a low calorie diet, but lower than those on a high calorie diet, at days 10, 20, and 30 (Figure 2H-J, Supplemental Table 1, 2).

Ultimately, this transient increase in mortality may be due to the effects of age, which could render flies less effective at dealing with the negative effects associated with a high calorie diet (20). Similar age-associated increases in mortality were observed in female flies maintained on a standard laboratory corn diet and mated at different ages, suggesting age-associated, rather than diet related, increases in mortality (21). Regardless, a previous history of a low calorie diet does not seem to be costly to young flies, and while switching may be acutely harmful to flies as they age, it does not impose a lifelong risk that is any higher than that of flies on a high calorie diet. *Taken together, shifting from a high to a low calorie diet is profoundly effective, even at an old age. Shifting from a low to a high calorie diet shows age-dependent increases in mortality*.

### Female egg production rapidly responds to caloric shifts

While diet has profound effects on the survivorship of male and female flies, females experience additional physiological changes due to the effects of diet on fecundity and egg production. Therefore, it is important to uncover how caloric shifts affect female egg production. Previous work from our group, and others, has demonstrated that female flies modify their egg production depending on diet (8,19,23). In order to carefully profile how egg production is modified when flies undergo caloric shifts, we measured egg production of females flies on high and low diets as well as after shifting diets in a manner similar to our lifespan studies; flies were shifted from high to low (and vice versa) calorie diets at two time points, 10 and 50 days. Numbers of eggs laid by each group were monitored daily and compared to the number of eggs produced by females shifted to the opposite diet and to eggs laid by female flies kept on a lifelong, low or high calorie diet. As expected, flies on a lifelong, low calorie diet produced fewer eggs than those on a high calorie diet (Figure 3A).

**Fig. 3.**
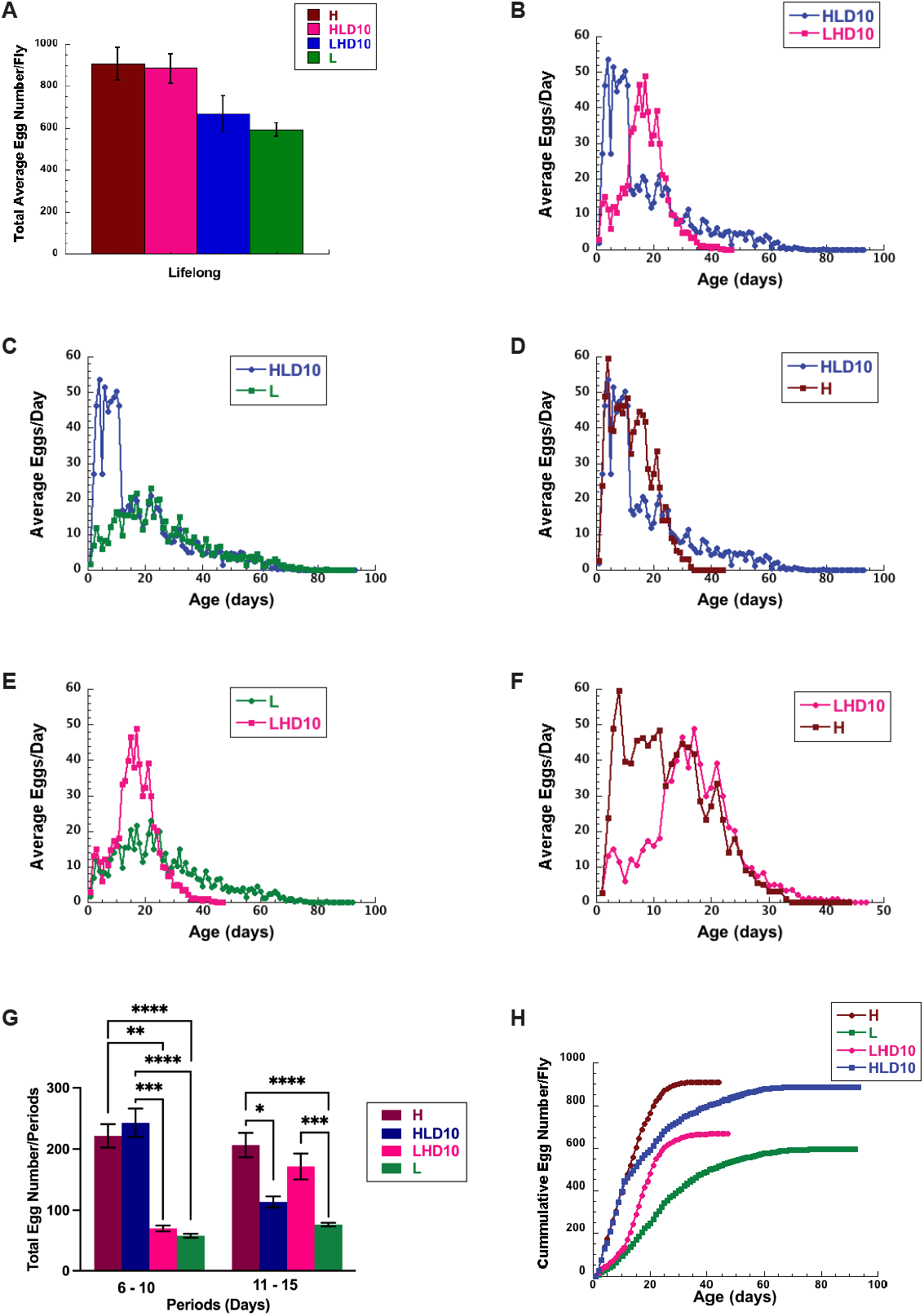
Calorie content of diet affects female life-long egg laying patterns. **A** Total average lifelong egg number produced by female *CS* flies kept on a high (brown), low (green), and flies shifted from a low to a high (magenta) or from a high to a low (blue) calorie diet on day 10. **B** Comparison of average egg production of female flies shifted from a low (L) to a high (H) or a high to low calorie diet at 10 days during whole life. **C** Comparisons of life long average daily egg production for female kept on a low (green) calorie diet to eggs produced by females shifted at 10 Days from a high to a low diet on day10 (blue). **D** Comparisons of life long average daily egg production for female kept on a high calorie diet (brown) to eggs produced by females shifted at 10 Days from a high to a low diet (blue). **E** Comparisons of life long average daily egg production for female kept on a low (green) calorie diet to eggs produced by females shifted at 10 Days from a low to a high calorie diet (magenta). **F** Comparisons of life long average daily egg production for female kept on a high (brown) calorie diet to eggs produced by females shifted at 10 Days from a low to a high calorie diet (magenta). **G** Total average egg production during period between 6-10 and 11-15 days produced by female *CS* flies kept on a high (brown), low (green), and flies shifted from a low to a high (magenta) or from a high to a low (blue) calorie diet on day 10. **H** Cumulative egg production in wild type female *CS* flies kept on a high (brown), low (green), and flies shifted from a low to a high (magenta) or from a high to a low (blue) calorie diet on day 10. Statistical analysis: The egg production data from 6-10 and 11-15 days were analyzed separately using the Kruskal-Wallis test. Post-hoc analysis was conducted using Dunn’s test, correcting for multiple comparisons using GraphPad Prism 9.4.1, Results represent SEM. There were 20 female flies in each experiment. p: 0.033 (*), 0.002 (**), 0.0002 (***), <0.0001 (****).

However, as with lifespan, flies were immediately responsive to dietary shifts, and would increase or decrease the number of eggs produced depending on diet (Figure 3B-F). Flies that were shifted from a high to low calorie diet at day 10 responded with a sudden drop in egg production to levels that matched flies on a lifelong, low calorie diet (Figure 3C). This was further illustrated by a similar total number of eggs laid from days 11-15 by females shifted from a high to low calorie diet on day 10 compared to females kept on a constant low diet during this same period (L=66, SEM=3.0, HLD10=114, SEM=9.0) (Figure 3G). Likewise, when shifted from a low to high calorie diet at day 10, the opposite was found to be true; flies responded by increasing the number of eggs produced (Figure 3E-F). Again, flies switched from a low to high calorie diet at day 10 produced similar numbers of eggs five days after shifting (days 11 – 15), to the number of eggs produced by females kept on a high diet from day 1 during the same period (H=206, SEM=19.8; LHD10=172, SEM=21.1) (Figure 3G, Table 3).

However, flies that were shifted from a low to high calorie diet did not produce the same cumulative number of eggs across their lifespan as flies raised on a constant, high calorie diet (Figure 3H). Since flies increased egg production to levels that match those on a high calorie diet immediately after shifting, this loss in cumulative production is likely due to the missed opportunity to produce a high number of eggs within the first 10 days of life, when females produce the highest number of eggs (21). Intriguingly, flies that were shifted from a high to low calorie diet at day 10 produced the same, cumulative number of eggs (∼900) across their lifespan as flies raised on a constant, high calorie diet (Figure 3A and H). This was strikingly different from flies that were maintained on a low calorie diet throughout their lifespan, which produced a much smaller number of eggs compared to flies on a high calorie diet (∼600 vs 900). This suggests that female flies can benefit from lifespan extension via CR without a sacrifice in lifelong fecundity, if dietary components or calories are provided early in life.

Surprisingly, female flies were still able to modify egg production in response to caloric shifts at day 50. When flies were shifted from a low to high calorie diet at this time point, there was a small increase in the number of eggs produced (Supplemental Figure 3A). Interestingly, a small but clear increase in egg production was noted when shifted to a high calorie diet despite very few flies remaining alive (Supplemental Figure 3B). For example, at day 58, approximately 5% of flies remained, but egg production maintained at about eight eggs per fly. This was starkly contrasted by flies switched to a low calorie diet, where about 80% of flies remained alive, but very few eggs, if any, were being produced. *Ultimately this finding highlights the fact that both fly lifespan and fecundity remain responsive to dietary shifts even at an old age, and that energy is immediately allocated to or from egg production depending on diet*.

### Metabolic alterations mediate the response to calorie restriction

Considering the dramatic response in lifespan and fecundity exhibited after late-life caloric shifts in female flies, we wanted to capture the underlying metabolic adaptations that may mediate such responses. To do this, we measured levels of triglycerides, glucose, glycogen, trehalose, and proteins in flies shifted from high to low (and vice versa) diets at two time points, 20 and 50 days.

The biggest change was observed in triglycerides levels, which showed an increasing trend when switched from a low to high calorie diet at day 20, which was significantly different from baseline (day 20, low) at day 25 (Figure 4A). This was exacerbated later in life, with triglyceride levels showing a significant increase each day the flies were on the high calorie diet (Figure 4A). The opposite was also true when flies were shifted from a high to a low calorie diet, levels of triglycerides began to drop. Again, this effect was more prominent in older flies, with a significant decline in triglyceride levels compared to baseline (day 50, low) (Figure 4A). *Together, this data suggests that female flies respond to dietary shifts by mobilizing or storing triglycerides, possibly to maintain tight regulation of glucose, trehalose, and glycogen levels*.

**Fig. 4.**
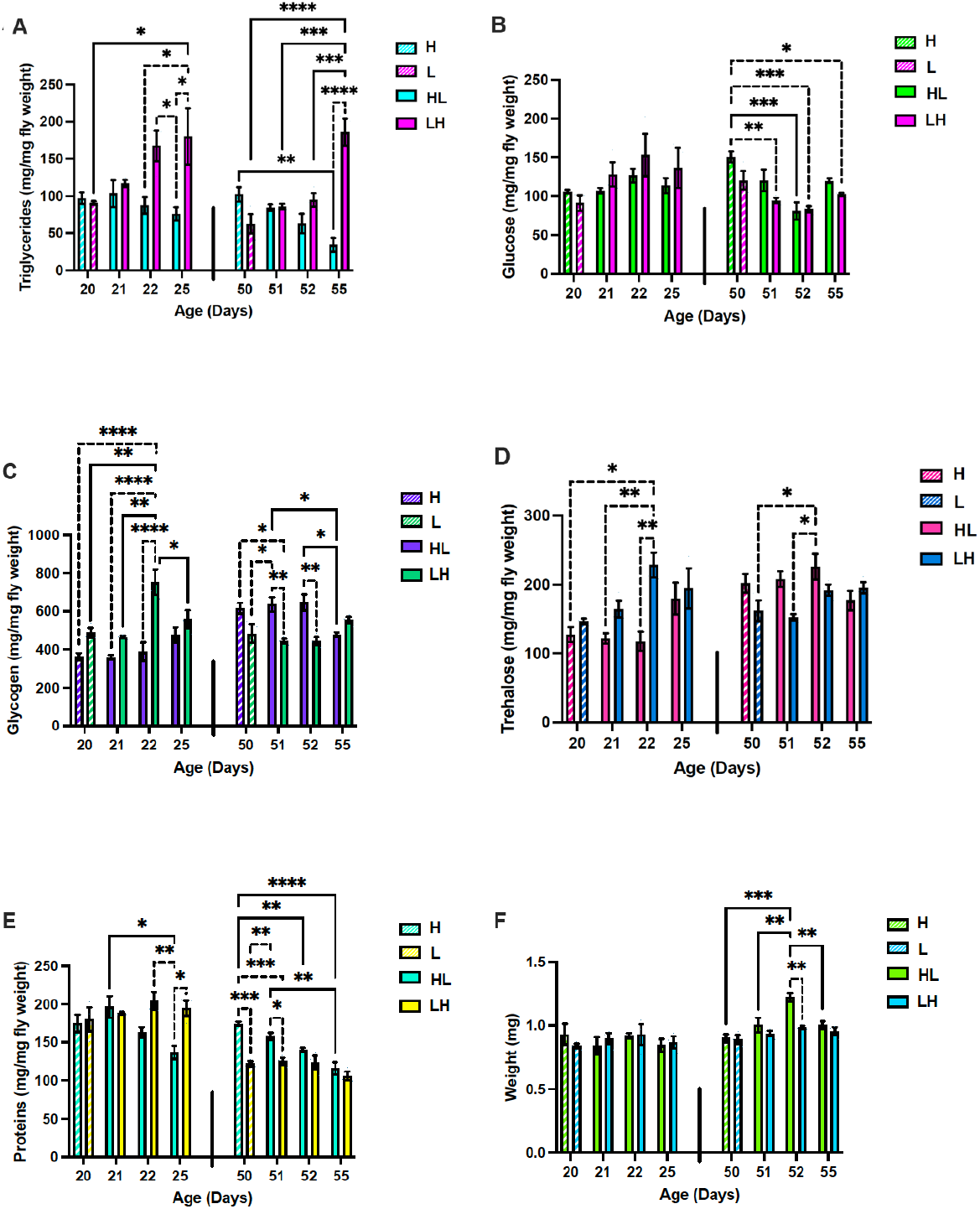
Effects of diet shifting on metabolism is age-dependent: Levels of triglycerides **(A)**, glucose **(B)**, glycogen **(C)**, trehalose **(D)**, protein **(E)**, and weight **(F)** in female flies on day 20 and 50 before shifting, and 1, 2, and 5 days after shifting from a high (H) to a low (L) calorie diet (HL) or L to H diet (LH). Error bars represent SEM. Data from 20 and 50 days were analyzed separately using two-way ANOVA. Post-hoc analysis was conducted using Tukey’s test, correcting for multiple comparisons. Results represent means ± SE of 3 biological replicates containing 10 flies per replicate. p: 0.033 (*), 0.002 (**), 0.0002 (***), <0.0001 (****).

Notably, flies appear to tightly regulate levels of glucose, glycogen, and trehalose at the 20 day time point, with levels remaining fairly stable upon both dietary switch regimes (Figure 4B-D). While flies shifted from a low to high calorie diet showed a significant increase in glycogen levels at day 22, the levels were not significantly different from baseline (day 20, low) at day 25, suggesting day 22 may be an anomaly (Figure 4C). Similarly, when switched from a high to low calorie diet at day 50, glycogen levels were significantly decreased by day 55, but there wasn’t otherwise a clear, daily, decreasing trend on this dietary regime (Figure 4C). However, levels of glucose did decrease in response to high to low shifts at day 50. Interestingly, protein levels were not responsive to high calorie diets, and remained stable when flies were shifted to a high calorie diet at every time point (Figure 4E). They were, however, responsive to low calorie diets, at both time points (Figure 4E). These results, paired with egg production, suggest flies may utilize the excess energy, particularly protein, associated with a high calorie diet to prioritize egg production. Once calories become sparing and protein levels begin to drop, this may cause a shift in resource allocation away from egg production. This has been demonstrated previously, as protein is a key dietary component required for egg production (19,24,25).

## Discussion

Together, our results demonstrate that *CS* female flies can benefit from late-life caloric shifts with no associated fitness cost and reveal the metabolic adaptations that mediate these events. Traditional evolutionary theory has suggested that caloric restriction is an environmental cue that signals to organisms the need to invest in somatic maintenance rather than reproduction (26). Under this paradigm, organisms would increase survival in order to have the opportunity to reproduce under more favorable conditions. However, this theory has been contested in studies which have identified dietary restriction to be associated with a hidden cost, which renders flies less fecund and more susceptible to death when returned to normal dietary conditions (14). However, this result may be dependent on strain, sex, diet, or more, as this was not the response identified in our and other studies (15). Here, we find female flies immediately respond to calorie restriction with a decrease in mortality when applied at any age tested. As expected, flies also decreased egg production; however, flies continued to produce eggs across their lifespan. Since flies live longer under these conditions, they ultimately produce the same cumulative number of eggs as flies on a high calorie diet, albeit at a slower rate. Further, when switched to a high calorie diet, younger flies did not experience an increase in mortality that was higher than flies raised on a constant, high calorie diet. These flies also responded with an immediate increase in egg production, which rose to match levels of flies on a high calorie diet. While flies switched from a low to a high calorie diet did not produce the same, cumulative number of eggs across their entire lifespan, likely due to missing out on the initial burst of production that occurs within the first ten days of life, starting on a low calorie diet did not hinder their ability to increase egg production when switched to a high calorie diet. Together, these results do not suggest a cost of calorie restriction, alternatively, they suggest the ability to optimize the fitness of female flies through the utilization of dietary shifts. For example, if flies are raised on a constant, low calorie diet, while they benefit from an increased lifespan, they produce significantly fewer eggs compared to flies on a constant, high calorie diet. However, if flies are first raised on a high calorie diet, and then switched to a low calorie diet, they can benefit from an extended lifespan with no cost to cumulative reproduction.

While older flies in our study displayed a transient increase in mortality when switched from a low to a high calorie diet, this increase did not endure as the flies aged. Additionally, flies in this dietary regime also responded with an increase in egg production. Similar to previous studies, this suggests that the increase in mortality when switched to a high calorie diet may be due to the cost of reproduction, mating, and increased vulnerability in aged flies (15,21,23). However, since flies in this group did not show a sustained increase in mortality after the switch occurred, and were able to begin producing similar numbers of eggs per fly as those in the high calorie group, our results suggest that age may render flies more vulnerable to the effects of reproduction and/or high calorie diets, but not that there is a fitness cost to calorie restriction.

This study also uncovered the metabolic adaptations that occur in response to dietary shifts. High to low shifting is associated with use of lipids, instead of glucose, as a major energy source. Similar metabolic adaptation was found in starved female flies when moved from satiated condition and was characterized by use of ketone bodies as a major source of energy instead of glucose (27). Flies may also prioritize maintaining sugar levels, possibly mobilizing or storing triglycerides in order to achieve homeostatic regulation of carbohydrates. Flies seem to be increasingly vulnerable to the effects of shifting at an older age, as those shifted to a high calorie diet show profound increases in triglyceride levels compared to baseline (day 50, high). This increase occurs within the context of elevated mortality, corroborating the pathological effects of high triglycerides on flies (28). It is tempting to think of this in the context of metabolic disease in humans, as high triglycerides are significantly associated with high risk of all-cause mortality (29). Further, our data are consistent with previous reports and further emphasize the importance of protein levels in egg production, as flies shifted to low calorie diets displayed decreasing levels of whole-body protein levels (24). This occurred within the context of egg production decline, suggesting flies are limited by protein to produce eggs. Counter to this, flies may maintain homeostatic protein levels through an increase in egg production, investing excess protein gained on a high calorie diet directly to egg production. Evidence has shown that limiting micronutrients, such as cholesterol, could also limit lifespan and egg production; therefore, we cannot discount the fact that flies on a high calorie diet were limited by other nutrients due to the increase in egg production (30,31). Future experiments which utilize our dietary shift regime, along with addition of nutrients like cholesterol, would be interesting to evaluate whether this is a limiting factor in our experimental conditions. However, as described before, cholesterol supplementation is not sufficient to fully compensate for the costs of a high protein diet, and therefore, is not expected to abrogate our results (15,30)

Ultimately, we find that female *CS* flies can benefit from CR, even when applied late in life. Flies can achieve substantial gains in lifespan despite very few flies remaining when shifted at the old age of 60 days. This gain in lifespan likely occurs by a decrease in risk of death, possibly mediated by metabolic adaptations (such as a decrease in triglycerides), a decrease in egg production, or a combination of other factors. Conversely, when switched to a high calorie diet, flies experience an increase in mortality, which appears to be more costly when flies are shifted at later ages. Despite this increase in mortality, flies are at no higher a risk than flies on a lifelong, high calorie diet, and have the capacity to produce the same number of eggs as their high calorie counterparts. This suggests that there is no cost associated with calorie restriction, which may be a sex specific finding. Finally, due to the dynamic ability of flies to respond to dietary shifts, our findings imply that fly fitness could be optimized, maximizing both lifespan and fecundity utilizing caloric shifts.

## Supporting information

Li et al., BioRxiv Supplemental Unformation

## Acknowledgments

We thank Dr. Gordon Carmichael for his helpful comments and suggestions. This work was supported by grants from the National Institute of Health: RO1AG059586, R01AG059586-03S1, the University of Connecticut (UConn) Claude D. Pepper Older Americans Independence Center (P30-AG067988) to B.R.; T32HG010463 to B.J.H. Rogina is a recipient of a Glenn Award for Research in Biological Mechanisms of Aging.

## References

1. Wilson KA, Chamoli M, Hilsabeck TA, Pandey M, Bansal S, Chawla G, et al. Evaluating the beneficial effects of dietary restrictions: A framework for precision nutrigeroscience. Cell Metabolism. 2021;33:2142–73.

2. Longo VD, Anderson RM. Nutrition, longevity and disease: From molecular mechanisms to interventions. Cell. 2022;185:1455–70.

3. Di Francesco A, Di Germanio C, Bernier M, de Cabo R. A time to fast. Science. 2018;362:770–5.

4. Rikke BA, Liao CY, McQueen MB, Nelson JF, Johnson TE. Genetic dissection of dietary restriction in mice supports the metabolic efficiency model of life extension. Experimental Gerontology. 2010;45:691–701.

5. Liao CY, Rikke BA, Johnson TE, Diaz V, Nelson JF. Genetic variation in the murine lifespan response to dietary restriction: from life extension to life shortening. Aging Cell. 2010;9:92–5.

6. Mitchell SJ, Madrigal-Matute J, Scheibye-Knudsen M, Fang E, Aon M, González-Reyes JA, et al. Effects of Sex, Strain, and Energy Intake on Hallmarks of Aging in Mice. Cell Metab. 2016;23:1093–112.

7. Kapahi P, Kaeberlein M, Hansen M. Dietary restriction and lifespan: Lessons from invertebrate models. Ageing Res Rev. 2017;39:3–14.

8. Bross TG, Rogina B, Helfand SL. Behavioral, physical, and demographic changes in Drosophila populations through dietary restriction. Aging Cell. 2005;4:309–17.

9. Campisi J, Kapahi P, Lithgow GJ, Melov S, Newman JC, Verdin E. From discoveries in ageing research to therapeutics for healthy ageing. Nature. 2019;571:183–92.

10. Partridge L, Pletcher SD, Mair W. Dietary restriction, mortality trajectories, risk and damage. Mechanisms of Ageing and Development. 2005;126:35–41.

11. Helfand SL, Rogina B. Genetics of Aging in the Fruit Fly, Drosophila melanogaster. Annu Rev Genet. 2003;37:329–48.

12. Helfand SL, Rogina B. Molecular genetics of aging in the fly: Is this the end of the beginning? Bioessays. 2003;25:134–41.

13. Mair W. Demography of Dietary Restriction and Death in Drosophila. Science. 2003;301:1731–3.

14. McCracken AW, Adams G, Hartshorne L, Tatar M, Simons MJP. The hidden costs of dietary restriction: Implications for its evolutionary and mechanistic origins. Sci Adv. 2020;6:eaay3047.

15. Sultanova Z, Ivimey-Cook ER, Chapman T, Maklakov AA. Fitness benefits of dietary restriction. Proc R Soc B. 2021;288:20211787.

16. Lee MB, Hill CM, Bitto A, Kaeberlein M. Antiaging diets: Separating fact from fiction. Science. 2021;374(6570):eabe7365.

17. Chao AM, Quigley KM, Wadden TA. Dietary interventions for obesity: clinical and mechanistic findings. J Clin Invest. 2021;131:e140065. 140065.

18. Woods JK, Kowalski S, Rogina B. Determination of the spontaneous locomotor activity in Drosophila melanogaster. J Vis Exp. 2014;(86).

19. Chapman T, Partridge L. Female fitness in Drosophila melanogaster: an interaction between the effect of nutrition and of encounter rate with males. Proc Biol Sci. 1996;263:755–9.

20. Li M, Macro J, Meadows K, Mishra D, Martin D, Olson S, et al. Late-life shift in caloric intake affects fly longevity and metabolism [Internet]. Physiology; 2023; http://biorxiv.org/lookup/doi/10.1101/2023.05.11.540262

21. Rogina B, Wolverton T, Bross TG, Chen K, Müller HG, Carey JR. Distinct biological epochs in the reproductive life of female Drosophila melanogaster. Mech. Ageing Dev. 2007;128:477–85.

22. Woods JK, Ziafazeli T, Rogina B. Rpd3 interacts with insulin signaling in Drosophila longevity extension. Aging (Albany NY). 2016;8:3028–44.

23. Rogina B. The effect of sex peptide and calorie intake on fecundity in female Drosophila melanogaster. ScientificWorldJournal. 2009;9:1178–89.

24. Skorupa DA, Dervisefendic A, Zwiener J, Pletcher SD. Dietary composition specifies consumption, obesity, and lifespan in Drosophila melanogaster. Aging Cell. 2008;7:478–90.

25. Min KJ, Tatar M. Drosophila diet restriction in practice: do flies consume fewer nutrients? Mech Ageing Dev. 2006;127:93–6.

26. Frankel S, Rogina B. Evolution, Chance, and Aging. Front Genet. 2021;12:733184.

27. Wilinski D, Winzeler J, Duren W, Persons JL, Holme KJ, Mosquera J, et al. Rapid metabolic shifts occur during the transition between hunger and satiety in Drosophila melanogaster. Nat Commun. 2019;10:4052.

28. Chatterjee N, Perrimon N. What fuels the fly: Energy metabolism in Drosophila and its application to the study of obesity and diabetes. Sci Adv. 2021;7:eabg4336.

29. Liu J, Zeng FF, Liu ZM, Zhang CX, Ling WH, Chen YM. Effects of blood triglycerides on cardiovascular and all-cause mortality: a systematic review and meta-analysis of 61 prospective studies. Lipids Health Dis. 2013;12:159.

30. Zanco B, Mirth CK, Sgrò CM, Piper MD. A dietary sterol trade-off determines lifespan responses to dietary restriction in Drosophila melanogaster females. eLife. 2021;10:e62335.

31. Zanco B, Rapley L, Johnstone JN, Dedman A, Mirth CK, Sgrò CM, et al. Drosophila melanogaster females prioritise dietary sterols for producing viable eggs. J Insect Physiol. 2023;144:104472.

