## Supplementary material for "Extended lifespan in female *Drosophila melanogaster* through late-life caloric restriction": Li et al., BioRxiv Supplemental Unformation

**Supplemental Figure Legends**

**Fig. S1 Shifting diets has immediate effects on female lifespan and mortality:** Survivorships (**A,C**) and mortality rates (**B,D**) between 50 and 85 (**A,B**) or 60 – 85 (**C,D**) of female flies shifted from a high (H) to a low (L) calorie diet at day 50 (HLD50) (**A,B**,) or day 60 (HDL60) (**C-D**) or from L to H diet on day 50 (LHD50) or day 60 (LHD60) (**H-L**). Number of flies: Number of flies: L=439, H=402, HLD50=217, HLD60=211, LHD50=226, LHD60=218. Survivorships curves and mortality rate were analyzed by long-rank test JMP16 program.

**Fig. S2 Shifting flies to diet with different calorie content affect female egg production:** Average daily egg production (**A**) and survivorships (**B**) between 50 and 60 days of female *CS* flies shifted from a low to a high calorie diet (LHD50) or from a high to a low (HLD50) calorie diet at day 50.

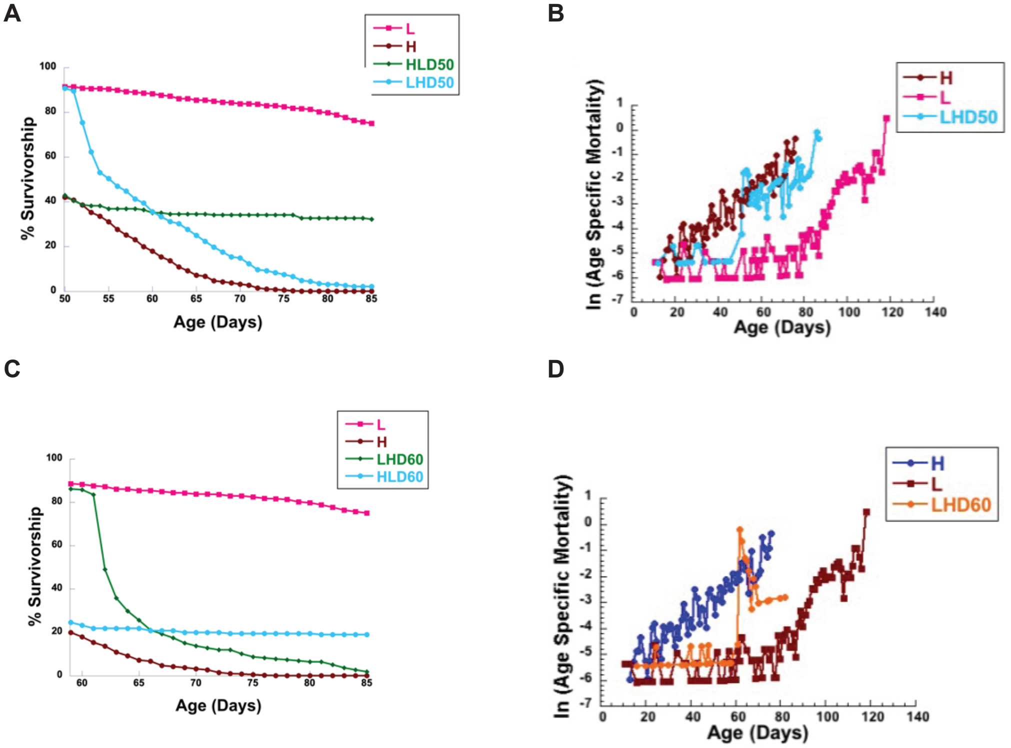


**Supplemental Figure 1**

A

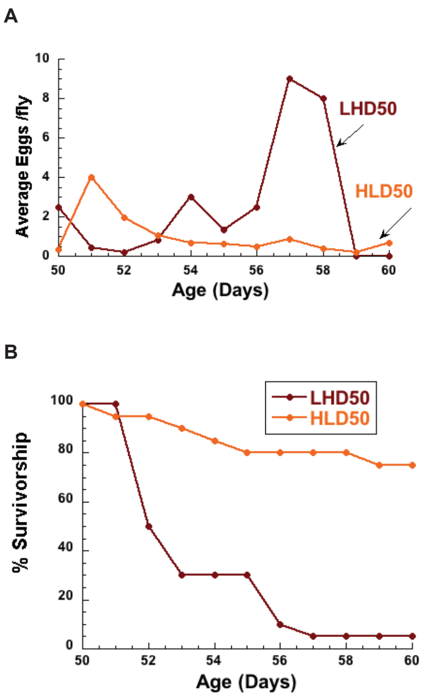


**Supplemental Figure 2**

**Supplemental Tables**

**Supplemental Table 1: Effects of shifting *Canton-S* female flies from a high (H) to a low (L) calorie diet on longevity compared to longevity of flies on lifelong high calorie diet.**

| Food | Time on L diet | N | Mean LS (% change) | $X^2$ | p | Maximal LS (% change) |
|---|---|---|---|---|---|---|
| H | Lifelong | 227 | 46.1 (-80.7) | | <0.0001* | 70.7 (-45.1) |
| L | Lifelong | 194 | 83.3 | | | 102.5 |
| H | Lifelong | 227 | 46.1 (-74.2) | 329.1587 | <0.0001* | 70.7 (-48.1) |
| HLD10 | D10 | 219 | 80.3 | | | 104.7 |
| H | Lifelong | 227 | 46.1 (-57.9) | 243.4023 | <0.0001* | 70.7 (-45.8) |
| HLD20 | D20 | 225 | 72.8 | | | 103.0 |
| H | Lifelong | 227 | 46.1 (-38.2) | 113.99962 | <0.0001* | 70.7 (-44.9) |
| HLD30 | D30 | 211 | 63.7 | | | 102.4 |
| H | Lifelong | 227 | 46.1 (-43.8) | 109.4052 | <0.0001* | 70.7 (-47.5) |
| HLD40 | D40 | 222 | 66.3 | | | 104.2 |
| H | Lifelong | 227 | 46.1 (-12.8) | 14.3933 | <0.0001* | 70.7 (-40) |
| HLD50 | D50 | 229 | 52.0 | | | 98.6 |

L=Low calorie diet
H=High calorie diet
HLD10= flies shifted from a high to low calorie diet at day 10
HLD20= flies shifted from a high to low calorie diet at day 20
HLD30= flies shifted from a high to low calorie diet at day 30
HLD40= flies shifted from a high to low calorie diet at day 40
HLD50= flies shifted from a high to low calorie diet at day 50
*Statistically significant

**Supplemental Table 2: Effects of shifting *Canton-S* female flies from a low (L) to a high (H) calorie diet on longevity in comparison to longevity of flies on lifelong low calorie diet.**

| Food | Time on L diet | N | Mean LS (% change) | $X^2$ | p | Maximal LS (% change) |
|---|---|---|---|---|---|---|
| L | Lifelong | 194 | 83.3 (44.6) | 359.1320 | <0.0001* | 102.5 (31.1) |
| H | Lifelong | 227 | 46.1 | | | 70.7 |
| L | Lifelong | 194 | 83.3 (40.6) | 258.5166 | <0.0001* | 102.5 (22.7) |
| LHD10 | D10 | 231 | 49.5 | | | 79.2 |
| L | Lifelong | 194 | 83.3 (40.2) | 286.1931 | <0.0001* | 102.5 (23.7) |
| LHD20 | D20 | 242 | 49.8 | | | 78.2 |
| L | Lifelong | 194 | 83.3 (34.8) | 248.8852 | <0.0001* | 102.5 (20.4) |
| LHD30 | D30 | 199 | 54.3 | | | 81.6 |
| L | Lifelong | 194 | 83.3 (36.5) | 313.1890 | <0.0001* | 102.5 (35.1) |
| LHD40 | D40 | 228 | 52.9 | | | 66.5 |
| L | Lifelong | 194 | 83.3 (32) | 325.2695 | <0.0001* | 102.5 (27.8) |
| LHD50 | D50 | 224 | 56.2 | | | 74.0 |

L=Low calorie diet
H=High calorie diet
LHD10= flies shifted from a high to low calorie diet at day 10
LHD20= flies shifted from a high to low calorie diet at day 20
LHD30= flies shifted from a high to low calorie diet at day 30
LHD40= flies shifted from a high to low calorie diet at day 40
LHD50= flies shifted from a high to low calorie diet at day 50
*Statistically significant
